# Streamlining Marker-less Allelic Replacement in *Streptococcus pneumoniae* Through a Single Transformation Step Strategy: easyJanus

**DOI:** 10.1101/2024.05.24.595743

**Authors:** Vipin Chembilikandy, Adonis D’Mello, Hervé Tettelin, Eriel Martínez, Carlos J. Orihuela

**Affiliations:** Department of Microbiology, Heersink School of Medicine, The University of Alabama at Birmingham, Birmingham, AL, USA; Department of Microbiology and Immunology, Institute for Genome Sciences, University of Maryland School of Medicine, Baltimore, MD, USA

**Keywords:** *Streptococcus pneumoniae*, Janus cassette, easyJanus, allelic exchange, transformation, homologous recombination, isogenic mutants, genetic deletion

## Abstract

The ability to genetically manipulate bacteria is a staple of modern molecular microbiology. Since the 2000’s, marker-less mutants of *Streptococcus pneumoniae* (*Spn*) have been made by allelic-exchange predominantly using the *kan*^*R*^*-rpsL* cassette known as “Janus”. The conventional Janus protocol involves two transformation steps using multiple PCR-assembled products containing the Janus cassette and the target gene’s flanking DNA. We present an innovative strategy to achieve marker-less allelic replacement through a single transformation step. Our approach involves the integration of an additional gene downstream region upstream of the Janus cassette, resulting in a modified genetic arrangement. This single modification reduced the number of required PCR fragments from five to four, lowered the number of assembly reactions from two to one, and simplified the transformation process to a single step. To validate the efficacy of our approach, we implemented this strategy to delete in *Spn* serotype 4 strain TIGR4 the virulence gene *pspA*, the entire capsular polysaccharide synthesis locus *cps4*, and to introduce a single nucleotide replacement into the chromosome. Notably, beyond streamlining the procedure, our method markedly reduced false positives typically encountered during negative selection with streptomycin when employing the traditional Janus protocol. Furthermore, and as consequence of reducing the amount of exogenous DNA required for construct synthesis, we show that our new method is amendable to the use of commercially available synthetic DNA for construct creation, further reducing the work needed to obtain a mutant. Our streamlined strategy, termed easyJanus, substantially expedites the genetic manipulation of *Spn* facilitating future research endeavors.

**IMPORTANCE:** We introduce a groundbreaking strategy aimed at streamlining the process for marker-less allelic replacement in *Streptococcus pneumoniae*, a Gram-positive bacterium and leading cause of pneumonia, meningitis, and ear infections. Our approach involves a modified genetic arrangement of the Janus cassette to facilitate self-excision during the segregation step. Since this new method reduces the amount of exogenous DNA required, it is highly amendable to the use of synthetic DNA for construction of the mutagenic construct. Our streamlined strategy, called easyJanus, offers significant time and cost savings, while concurrently enhancing the efficiency of obtaining marker-less allelic replacement in *S. pneumoniae*.

## INTRODUCTION

*Streptococcus pneumoniae*, an opportunistic pathogen, is a leading cause of otitis media, community-acquired pneumonia, bacteremia, and meningitis (1, 2). First described by Pasteur and Sternberg in 1881 (3, 4), work with the pneumococcus by Avery, McLeod, and McCarty in 1944 demonstrated that DNA was the molecule responsible for the transfer of heritable traits and ushered in the modern era of molecular biology (5). In 2001, the first annotated genome of *S. pneumoniae* was published (6). Since then, *S. pneumoniae* has become one of the most sequenced bacteria in the world, with >40,000 genomes being publicly available (7). This wealth of genomic data has empowered researchers to identify genome regions encoding potential virulence determinants, moreover, better understand how this pathobiont is able to cause disease (8, 9). Along such lines, the genetic manipulation of *S. pneumoniae*, i.e., the creation of genetic mutants, has played a pivotal role in unraveling its physiology and the basis for pathogenesis.

*S. pneumoniae*’s ability for natural competence has been and continues to be a boon for its genomic manipulation. Investigators have learned to co-opt this process and easily replace DNA on the genome using mutagenic constructs that target a gene based on flanking DNA sequences that serve as the sites for homologous recombination. Suicide vector plasmids and PCR-generated mutagenic constructs have been used to delete individual genes, operons, pathogenicity islands, and even introduce single-nucleotide replacements (10, 11). Critically, the principal molecular method used today to create marker-less mutants in *S. pneumoniae*, as well as many other bacterial species is the Janus cassette (12). This DNA construct, flanked by the upstream and downstream regions of the target locus, features a kanamycin resistance marker and a counter selectable *rpsL* gene which confers dominant streptomycin sensitivity in a resistant background. After kanamycin selection, which results in selection of mutants lacking the gene of interest, the second transformation step introduces the allele of choice flanked by the same DNA regions. Streptomycin resistance as result of *rpsL* is in turn restored, and this is used to select for the loss of Janus and the acquisition of the desired allele including deletion mutants. Importantly, the use of streptomycin selection has posed challenges due to the high frequency of false positives arising from spontaneous mutations in *rpsL*^*+*^ in Janus, necessitating a tedious secondary screen. To address this, a modified Janus cassette, termed Sweet Janus, was introduced, incorporating an additional counter-selectable marker, *sacB* (13). While enhancing negative selection efficiency, it still required two transformation steps and selection in the presence of sucrose adding complexity to the procedure.

In this context, we present a novel single-step transformation methodology for allelic replacement that significantly reduces time and costs. What is more, our approach can take full advantage of emergent capabilities in synthetic DNA construction, precluding the requirement of PCR to generate the mutagenic construct, further saving time and costs. By simplifying and enhancing the efficiency of genetic manipulation, our approach holds immense promise for accelerating research and advancing our understanding of this significant bacterial pathogen.

## MATERIALS AND METHODS

### Bacterial strain and culture conditions

S. *pneumoniae* GPSC27, serotype 4, strain TIGR4 *rpsL*^*+*^ was used for the study (6) . Bacteria were routinely grown in Todd-Hewitt broth supplemented with 0.5% yeast extract (THY) or on tryptic soy agar (5% sheep blood) plates at 37°C with 5% CO_2_. Antibiotics were added, when necessary, at the following concentrations: kanamycin at 200 μg/mL and streptomycin at 200 μg/mL.

### Construction of DNA fragments

Oligonucleotides used in this study are listed in Table S1. DNA fragments were generated using specific sets of primers designed to amplify the Janus cassette, along with different-sized upstream and downstream DNA regions. Amplification proceeded for 36 cycles as follows: 20 s at 94°C, 30 s at 54°C, and 1.45 min at 72°C, followed by a 5-min extension cycle using Q5^®^ High-Fidelity 2X Master Mix (New England Biolabs). PCR products were then purified using QIAquick® Gel Extraction Kit (Qiagen, Valencia, CA). Of note, we added an additional 25 bp to the second downstream fragment so that this fragment contains a primer binding site that is absent in the middle downstream fragment. This allows the entire construct to be PCR amplified after their assembly if the frequency of mutant generation following transformation with the assembled DNA construct product was low and needed repeating. Custom ordered synthetic DNA fragments were obtained from Twist Biosciences Inc. (South San Francisco, CA). PCR / synthetic DNA fragments were assembled using assembly master mix (NEBuilder® HiFi DNA Assembly).

### Natural transformation of *Spn*

TIGR4 was grown in THY media until the culture reached an optical density at OD_621_ of ∼0.2. Subsequently, 200μL were transferred to competent media (5mL THY, 50μL of 20% glucose, 250μL of 5% BSA, 10μL of 10% CaCl_2_ and). Competence was induced with 1μL of CSP 2 (Eurogentec) and then incubated at 5% CO_2_ and 37°C for 10 minutes. Subsequently 500 μL of competent bacteria were added to microcentrifuge tubes containing 20 μL assembled mutagenic DNA constructs and incubated for 30 minutes in 5% CO_2_ and 37°C. Following incubation, the bacteria were transferred to 7mL of THY and grown for 3-4 hours. For the selection of transformant colonies, 100μL of culture was spread on a blood agar media with kanamycin (200μg/mL). For the segregation step, kanamycin resistant colonies were grown in 3mL of THY for roughly 3-4 hours. Subsequently, 100μL of culture was spread on a plate with streptomycin (200μg/mL). Mutants were confirmed by PCR followed by sequencing.

## RESULTS

Employing the Janus cassette for allelic replacement involves two steps of transformation. First with a donor DNA fragment containing both upstream and downstream regions of the target gene flanking the Janus cassette, and subsequently with the fragment containing the desired allele between the upstream and downstream regions; for the purpose of gene deletion, no DNA is present between the corresponding flanking regions (Fig. 1A). Our hypothesis posited that incorporating an additional downstream sequence of the target gene upstream of Janus would result in the self-excision of the Janus cassette post-integration and first selection, obviating the need for a subsequent transformation step (Fig 1B). To validate this hypothesis, we aimed to delete the *pspA* gene using this approach. Initially, our construct featured a configuration of near equal length upstream DNA (Up), downstream DNA (Down), Janus, and downstream DNA (500bp-Up:500bp-Down:Janus:525bp-Down). Note that the downstream fragment included a 25bp primer binding site at its 3’ end for purposes of PCR amplification (see Methods). Transformation of this construct yielded kanamycin-resistant colonies, and subsequent PCR analysis revealed that 55% of the colonies corresponded to allelic replacement of *pspA* by the Janus cassette (Table 1), while the remaining 45% resulted from Janus integration downstream of the gene (Fig. S1). Following PCR validation for the correct co-integrant, we allowed self-segregation of Janus by cultivating independent colonies on broth THY, followed by selection of segregants through streptomycin selection. PCR analysis demonstrated that 92% of the streptomycin-resistant colonies harbored the designed deletion of the *pspA* gene. The low frequency of false positives during this step indicated that the high frequency of intramolecular homologous recombination between the duplicated downstream region of our construct surpassed the frequency of *rpsL* revertants.

**Table 1.** Frequency of successful integration and segregation using easyJanus during construction of *ΔpspA, Δcps4* and Sp_0837 529 V→E.

|  | Donor DNA | DNA alignment* | Frequency (%) |  |
| --- | --- | --- | --- | --- |
|  |  |  | Integration | Segregation |
| *pspA* | Assembled-PCR | 500bp: **500bp**: Janus: **525bp** | 55 | 91.66 |
| *pspA* | Assembled-PCR | 1000bp: **500bp**: Janus: **525bp** | 58.33 | 98.88 |
| *pspA* | Synthetic | 1000bp: **500bp**: Janus: **525bp** | 82.10 | 99 |
| *pspA* | Synthetic | 500bp: **250bp**: Janus: **275bp** | 75.79 | 96 |
| *cps* | Synthetic | 500bp: **250bp**: Janus: **275bp** | 100 | 90.66 |
| V→E | Synthetic | 500bp: **250bp**: Janus: **275bp** | 100 | 86.66 |

\*Bold denotes sections with identical DNA sequence, except for 25 the additional bases included for amplification primer design.

**Figure 1.**
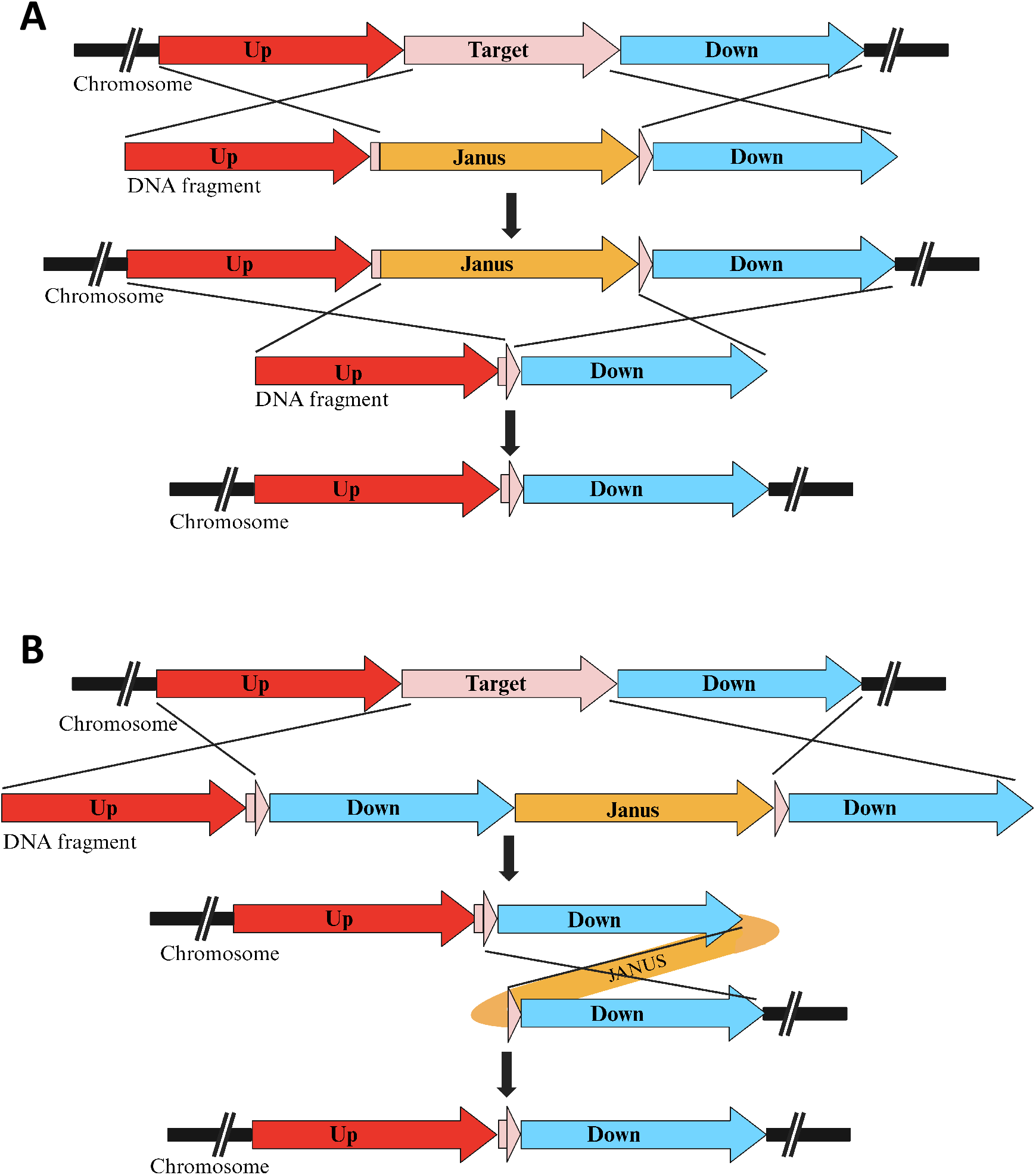
Schematic comparison between **A)** the conventional method for allelic replacement utilizing the Janus cassette and **B)** our novel easyJanus approach. In the traditional method, two donor DNA fragments and two transformation steps are necessary. Conversely, the new strategy involves a single DNA fragment as a donor, with only one transformation step required, facilitated by the self-segregation event through intramolecular homologous recombination.

To enhance the frequency of obtaining the allelic replacement of the gene instead of downstream integration of Janus in the initial step, we augmented the proportion of the upstream region of the target gene in our construct compared to downstream regions, i.e., 500bp-Up:250bp-Down:Janus:275bp-Down (Fig. S2). By providing more upstream region than downstream, we intended to augment the likelihood of the first crossover occurring within the 500bp upstream region of the target gene. As anticipated, this led to a substantial increase in the frequency of obtaining the replacement of *pspA* by Janus from 55% to 82% in the first transformation step. Once again, following selection of the allelic replacement, segregation resulted in 100% of streptomycin-resistant colonies harboring a deletion of the *pspA* gene (Table 1).

The opportunity for a single DNA donor to achieve marker-less allelic replacement prompted us to explore the use of synthetic DNA as a donor instead of assembled PCR products. Due to limitations in synthesizing DNA molecules containing duplicated sequences, we divided our construct into two fragments for synthesis that harbored overlapping regions for subsequent HiFi assembly. In this instance, our synthetic DNA donor, comprising upstream DNA, downstream DNA, Janus, downstream DNA (1000bp-Up:500bp-Down:Janus:525bp-Down), generated allelic replacement of *pspA* with a frequency of 82%, similar to that of the PCR-assembled counterpart, and 99% during segregation (Table 1). Subsequently, to reduce the cost of the synthetic DNA construct, while maintaining the frequency of allelic replacement, we reduced the construct to half its size while maintaining the same flanking segment order and relative proportion, i.e., 500bp:250bp:Janus:275bp. Employing this construct yielded a similar frequency of allelic replacement (75.79%), with 96% during segregation (Table 1).

Given that allelic exchange often entails not only gene deletion but also the removal of larger DNA segments or single-nucleotide substitutions, we applied our strategy to delete the entire *cps4* locus, encompassing 25 genes spanning 21,592bp. Using synthetic fragments, we successfully deleted *cps4*, achieving a 100% efficacy in the first step and 90.66% counterselection (Table 1). Validation of the *cps4* deletion was performed through PCR followed by sequencing. We also evaluated the efficiency of inserting point mutations into gene SP_0837, altering the nucleotides TT to AA and thereby changing the encoded amino acid from 529-VRIEL-533 to 529-ERIEL-533. We achieved this mutagenesis using synthetic fragments with a frequency of 100% in the first step and 86.66% during counterselection (Table 1).

Finally, we felt that a more automated approach for the creation of mutagenic donor constructs should be available. To this end, we wrote a Perl script that takes a whole genome sequence (or any sequence spanning regions targeted for modification and their flanks) as input, along with a four-column tab-delimited list of targets in the form of contig identifier, target name, start coordinate, and end coordinate of the target on the input sequence provided. As an example, to generate constructs to delete *pspA*, the input can be the TIGR4 whole genome sequence (NCBI NC_003028) then the target list containing one line can be: “NC_003028 SP_0117 118423 120657”. The script then extracts relevant sequences from the genome to generate two constructs of 1,199bp each: Fragment_1 made of 500bp-Up:250bp-Down:449bp-Janus, and Fragment_2 made of 924bp-Janus:275bp-Down. The two pieces of Janus overlap by 40bp for i*n vitro* assembly to create a single fragment prior to transformation. For portability, the script has been embedded in a Docker image – https://hub.docker.com/r/admellodocker/easyjanus_design – that can be installed on a desktop or laptop running Windows, MacOS, or Linux.

## DISCUSSION

The successful implementation of allelic replacement strategies, particularly in bacterial genetics, holds significant implications for understanding gene function, microbial pathogenesis, and genetic engineering applications. In this study, we introduce a new strategy aimed to streamline allelic replacement in *Spn* using the Janus cassette. Our study demonstrates the efficiency of this new methodology, thus paving the way for easier and more rapid genetic manipulations in *Spn* and other bacteria. Briefly, our streamlined methodology also allows for quick oligonucleotide sequence design for transformation through our portable Docker-Perl script, reduces the number of required PCR products (if used) from five to four, consolidates assembly reactions from two to one, and crucially reduces the need for two transformation steps down to one (Table 2). Moreover, the easyJanus protocol significantly slashes the time needed to generate mutants from the conventional five day process to as few as two days.

**Table 2.**
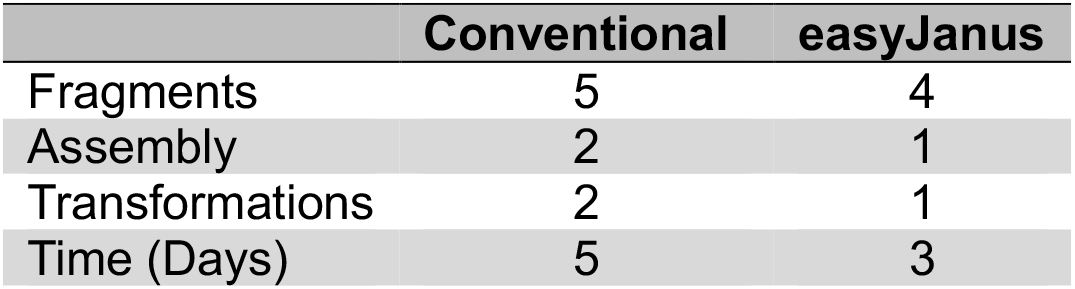
Comparison between the traditional protocol using the Janus cassette and the easyJanus system.

|  | Conventional | easyJanus |
| --- | --- | --- |
| Fragments | 5 | 4 |
| Assembly | 2 | 1 |
| Transformations | 2 | 1 |
| Time (Days) | 5 | 3 |

Our initial experiments revealed the presence of undesired integrant stemming from a double crossover event between the two downstream regions of the target gene flanking the Janus cassette and the chromosome’s downstream region (Fig. S1). Consequently, our subsequent effort focused on optimizing the design of the donor DNA fragment to preferentially facilitate efficient allelic replacement. By strategically adjusting the size of the upstream and downstream regions surrounding the target gene, we successfully increased the frequency of obtaining accurate integrants and reduced the size of the construct such that the use of synthetic DNA was more feasible. It is our opinion that the use of synthetic DNA to create the donor construct is highly advantageous due to its clear advantages in terms of scalability, reproducibility, cost effectiveness, and sequence customization. Synthesized DNA constructs generated allelic replacements at a frequency comparable to that achieved using PCR-assembled donors. This underscores the feasibility and efficiency of utilizing synthetic DNA constructs in allelic replacement.

We confirmed the application of our methodology beyond single gene deletions to encompass larger genetic modifications, such as the removal of entire gene clusters or the introduction of specific point mutations. Our successful deletion of the *cps* locus, comprising multiple genes spanning a considerable genomic region, underscores the robustness and scalability of our approach for manipulating complex genetic loci. Additionally, our ability to introduce precise point mutations in target genes highlights the versatility and precision afforded by our methodology, thereby enabling fine-tuned genetic modifications with broad implications for functional genomics and molecular biology studies. Based on our findings, we propose a refined protocol: initially selecting five kanamycin-resistant colonies followed by the segregation of each selected colony to identify two streptomycin-resistant colonies, resulting in a total of ten colonies (Fig. 2). Our analysis indicates that this approach yields the desired mutant with a probability exceeding 98%. Notably, our strategy offers opportunities for enhancement. For instance, our data reveal that increasing the proportion of the upstream region significantly enhances the likelihood of achieving the desired allelic replacement. Furthermore, optimizing the fragment size can potentially reduce the cost of DNA synthesis, particularly when utilizing synthetic DNA.

**Figure 2.**
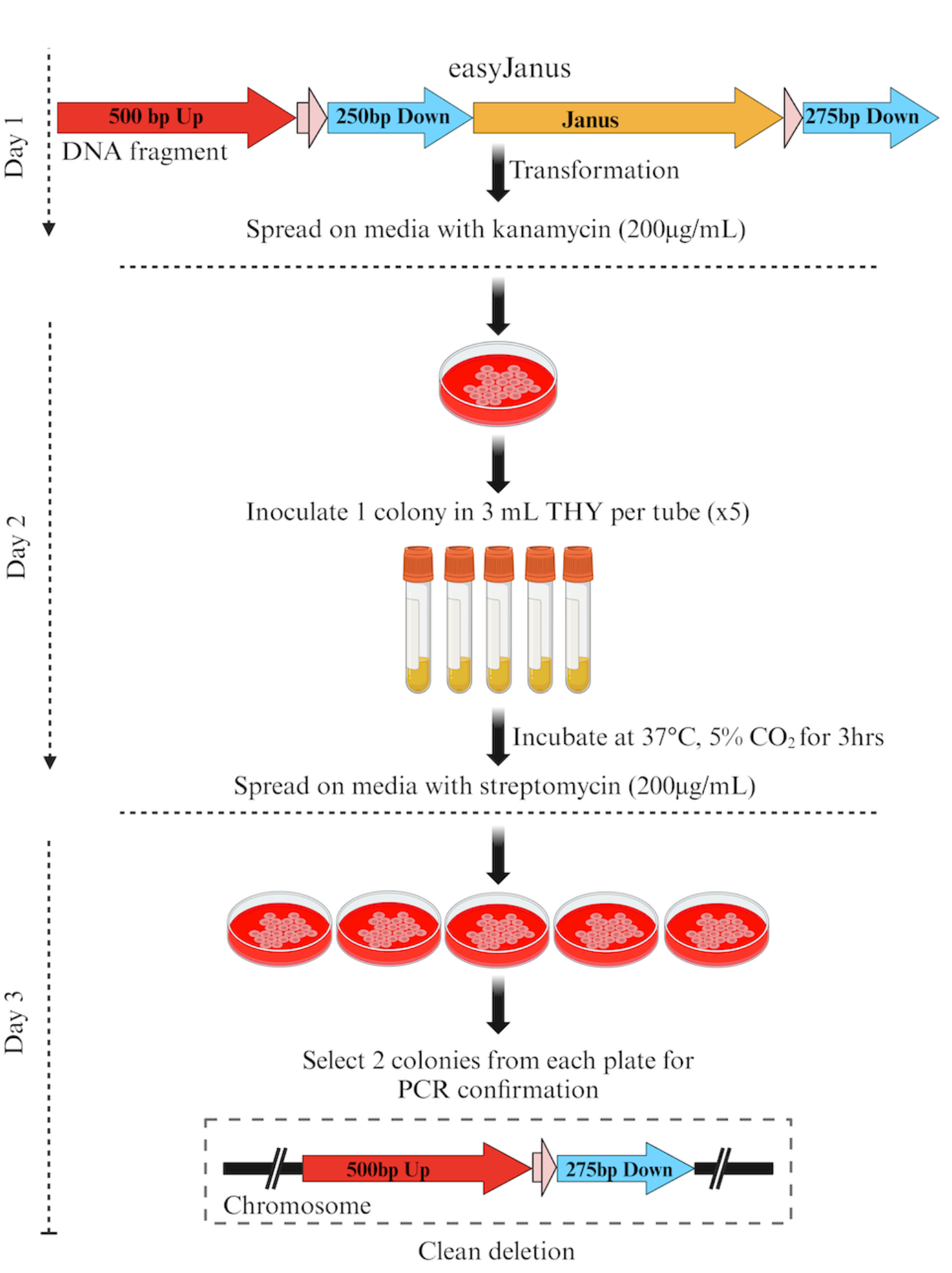
Schematic representation of the proposed 3-day protocol for achieving allelic replacement in *Spn*. We recommend selecting 5 kanamycin-resistant colonies, followed by the segregation of each selected colony to identify 2 streptomycin-resistant colonies. Our thorough analysis affirms that this method offers a probability of over 98% in obtaining the desired mutant.

Overall, our study demonstrates the efficacy and versatility of the proposed allelic replacement approach in facilitating precise and marker-less genetic manipulations in bacterial systems. For this reason, we have termed this approach easyJanus. By systematically optimizing donor DNA design and leveraging synthetic DNA technologies, we hope to have streamlined the process of genetic engineering for other investigators, opening up new avenues for studying gene function, microbial physiology, and biotechnological applications. It is for this reason that we developed a computer script that automates the generation of sequences for two 1,199bp fragments to be synthesized and directly used for mutant generation. As indicated, the script runs in a portable and easy to use Docker application that can be deployed on any of the three most common operating systems. We note that sequence coordinates provided to the script can be for any short or long region within the genome, and could, for instance, encompass multiple genes such as the *cps* capsule synthesis locus.

In ending, we feel this breakthrough, alongside strides in synthetic DNA technology, now renders it plausible, and within reasonable cost, to assemble a comprehensive library of markerless tagged mutants for *Spn*. Such a library holds the promise of surpassing existing mutant collections based on insertional disruption of the target gene in random fashion, offering on-demand isogenic mutants without inducing polar effects on neighboring genes.

---

**Figure S1.**
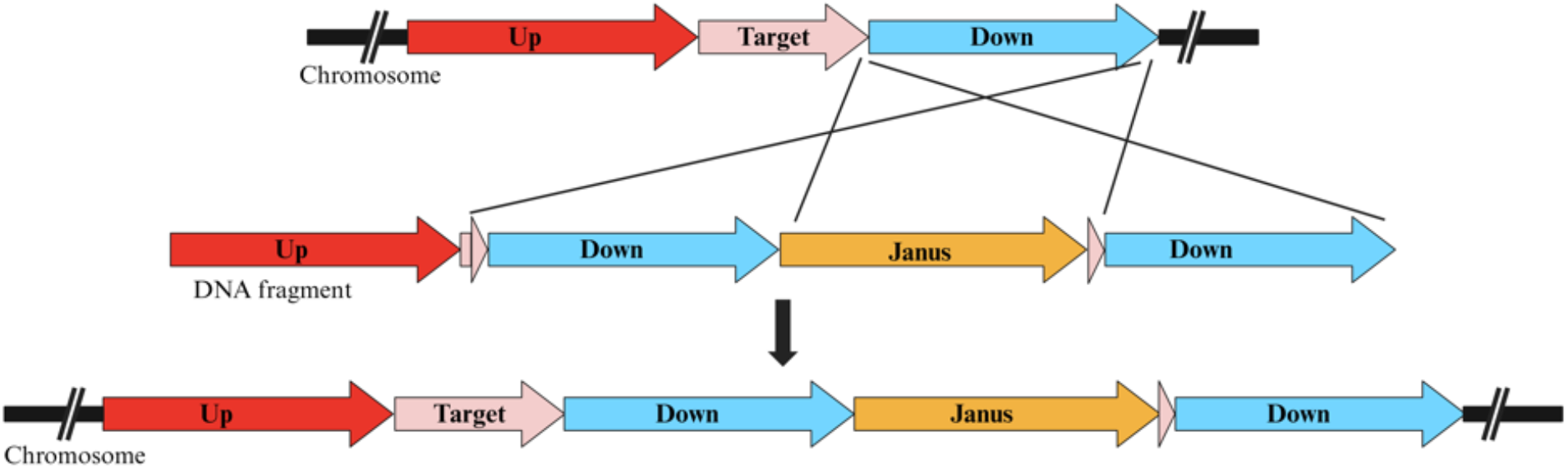
Double recombination events occurring between the downstream region of the target gene and the duplicated downstream region within the donor fragment result in Janus integration downstream of the gene.

**Figure S2.**
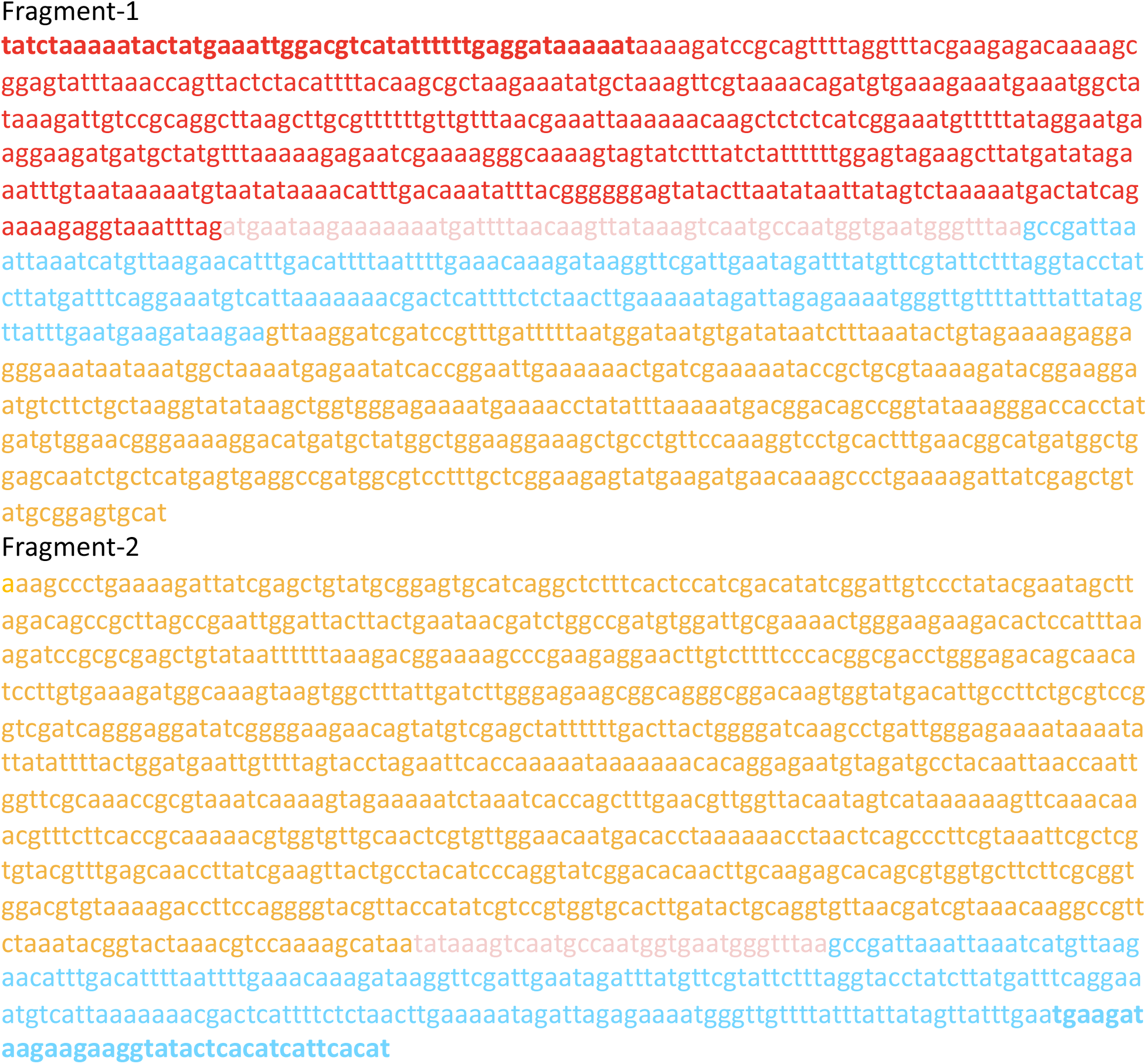
Sequences of the designed fragments for the deletion of the *pspA* gene. The red region indicates the upstream region of the *pspA* gene. The pink regions represent the remaining parts of the *pspA* gene after deletion. The blue regions are the downstream segments of the *pspA* gene. PCR primer binding sites, highlighted with bold red letters for the forward primer and bold blue letters for the reverse primer, are designed to amplify the assembled product and/or confirm proper allelic replacement. The color coding in this figure is consistent with Figure 1B for ease of reference.

**Table S1.**
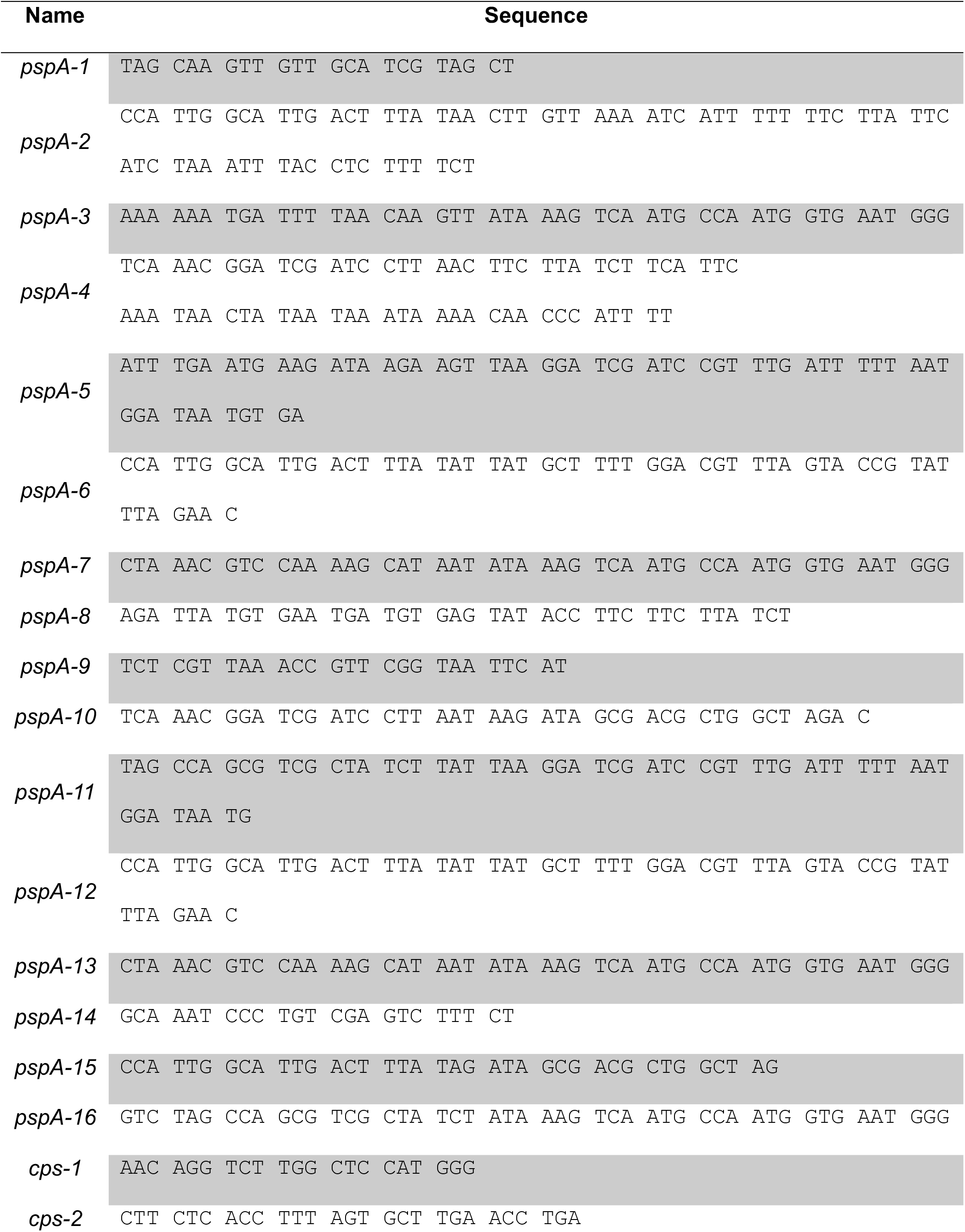
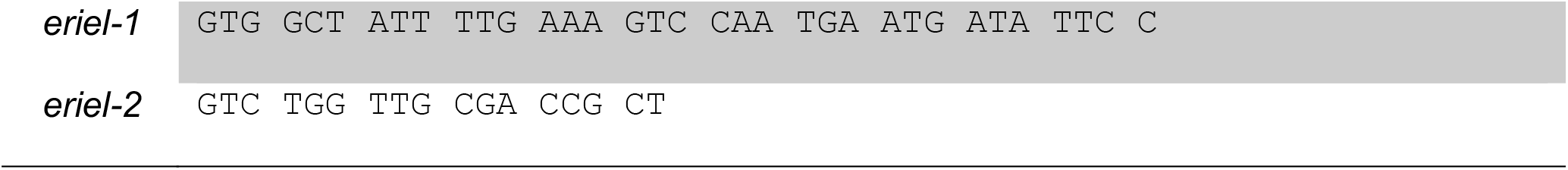
Oligonucleotides used in this study.

